# Variable penetrance of the 15q11.2 BP1–BP2 microduplication in a family with cognitive and language impairment

**DOI:** 10.1101/095919

**Authors:** Antonio Benítez-Burraco, Montserrat Barcos-Martínez, Isabel Espejo-Portero, M^a^ Salud Jiménez-Romero

## Abstract

The 15q11.2 BP1–BP2 region is found duplicated or deleted in people with cognitive, language, and behavioral impairment. *Case presentation*. We report on a family (the father and three male twin siblings) who presents with a duplication of the 15q11.2 BP1-BP2 region and a variable phenotype: whereas the father and the fraternal twin are normal carriers, the monozygotic twins exhibit severe language and cognitive delay and behavioral disturbances. The genes located within the duplicated region are involved in brain development and function, and some of them are related to language processing. *Conclusions*. The probands’ phenotype may result from changes in the expression level of some of these genes important for cognitive development.

## INTRODUCTION

Rare or sporadic conditions involving language deficits and resulting from chromosomal rearrangement or copy-number variation provide with crucial evidence of the genetic underpinnings of the human faculty of language. The BP1-BP2 region at 15q11.2 is commonly found duplicated or deleted in patients referred for microarray analysis. Copy number variation (CNV) of this region usually results in developmental delay, intellectual disability, speech and language delay, behavioural disturbance, and motor delay, but also in neuropsychiatric conditions like autism, schizophrenia; dysmorphic features and epilepsy are also reported (Burnside et al. 2011, Abdelmoity et al. 2012, Cafferkey et al. 2014, Cox and Butler 2015; Picinelli et al. 2016). Changes in the BP1-BP2 region have been claimed to increase the predisposition to language delay in Type I deletion subjects with Angelman syndrome and Prader-Willi syndrome (Burnside et al., 2011), two conditions resulting from the deletion of the BP1-BP3 (type I) or the BP2-BP3 (type II) regions at 15q11.2. Deletions and duplications of the BP1-BP2 region are equally common, but apparently, deletions impact more than duplications on cognitive development and language acquisition (Burnside et al. 2011). At the same time, variable penetrance of this CNV is commonly reported (see Hashemi et al. 2015 and Vanlerberghe et al.2015 for discussion). As noted by many authors (e.g. De Wolf et al. 2013) studies on how the deletion or the duplication of this region is transmitted over generations may help to understand this heterogeneity. Likewise, we still lack an in-depth account of language deficits and language development in people affected by CNV of the BP1-BP2 region.

In this paper, we report on three twin children who bear a duplication of the BP1-BP2 region transmitted by their father, providing a detailed characterization of their linguistic phenotype. Interestingly, the two monozygotic twins present with severe language delay, motor problems, and behavioural disturbances, including autistic features, whereas the fraternal twin and the father show a milder phenotype with no major impairment.

## METHODS

### Cognitive, linguistic, and behavioral evaluation

Global development of the three children was assessed with the Spanish version of the Battelle Developmental Inventories (De la Cruz & González, 2011). This test comprises 341 items and includes specific subtests for evaluating receptive and expressive communication skills, gross and fine motor skills, cognitive development, personal/social development, and adaptive abilities.

Autistic features in the two monozygotic twins were assessed with the Spanish version of the Modified Checklist for Autism in Toddlers (M-CHAT) (Robins et al. 2011). M-CHAT is a screening (not a diagnosing) tool for toddlers between 18 and 60 months. It comprises 23 questions. A positive score is indicative of the possibility of suffering from the disease.

Psycholinguistic development of the fraternal twin was assessed in more detail with the Spanish version of the Illinois Test of Psycholinguistic Abilities (ITPA) (Ballesteros & Cordero, 2004). This test is aimed to evaluate the child’s abilities in two different channels of communication (auditive-vocal and visuo-motor), two levels of organization (automatic and representative), and three psycholinguistic processes (receptive, associative, and expressive). Items are organized in 13 subtests: Auditory reception (AR); Visual reception (VR), Auditory Sequential Memory (ASM); Visual Sequential Memory (VSM); Auditory Association (AA); Visual association (VA); Auditory Closure (AC); Visual Closure (VC); Manual expression (ME); Verbal expression (VE); Grammatical Closure (GC); and Sounds Combination (SC). Because of the severity of their cognitive impairment, this test could not be applied to the monozygotic twins.

The verbal abilities of the fraternal twin were also assessed with the verbal component of the Spanish version of WISC-IV (Wechsler, 2005), whereas the Spanish version of the WAIS-III (Wechsler, 1999) was used to evaluate the parents’ verbal performance. This component of the test comprises 6 tasks aimed to evaluate two different domains of language processing: verbal comprehension (Similarities (S), Vocabulary (V), Information (I), and Comprehension (CO)) and working memory (Digits (D) and Arithmetic (A)).

### Molecular cytogenetic analysis

#### Karyotype analysis

Peripheral venous blood lymphocytes were grown following standard protocols and collected after 72 hours. A moderate resolution G-banding (550 bands) karyotyping by trypsin (Gibco 1x trypsin^®^ and Leishmann stain) was subsequently performed. Microscopic analysis was performed with a Nikon^®^ eclipse 50i optical microscope and the IKAROS Karyotyping System (MetaSystem^®^ software).

#### Angelman syndrome determinants

Fluorescent in situ hybridization (FISH) with the LSI Prader-Willi/Angelman probes (15q11-q13 SNPRN and 15q11-q13 D15S10 Izasa^®^) was conducted to detect microdeletions of the 15q11-q13 locus and/or of *UBE3A,* located at 15q11.2. Metaphase spreads were harvested from peripheral blood as above. Slides were analyzed with a Nikon^®^ eclipse 50i optical microscope and the Metasystem ISIS^®^ software.

#### Multiplex ligation-dependent probe amplification (MLPA)

MLPA was performed to detect abnormal CNV of the subtelomeric regions of the probands’ chromosomes. The SALSA MLPA kits P036 and P070 from MRC-Holland were used according to the instructions provided by the supplier.

#### Microrrays for CNVs search and chromosome aberrations analysis

DNA samples (total amount: 500 ng) extracted from blood were hybridized on a CGH platform (Cytochip Oligo ISCA 60K).The DLRS value was > 0.10. The platform included 60.000 probes. Data were analyzed with Agilent Genomic Workbench 7.0 and the ADM-2 algorithm (threshold = 6.0; aberrant regions had more than 5 consecutive probes).

## RESULTS

### Clinical History

The probands are three twin brothers (one fraternal twin [hereafter FT] and two monozygotic twins [hereafter MT1 and MT2]) resulting from in vitro fertilization and born after 35 weeks of normal gestation by Cesarean birth to a healthy 31 years old female. The most relevant clinical data at birth are summarized in table 1

**Table 1.** Summary table with the most relevant clinical findings at birth.

|  | **MT1** |  | **MT2** |  | **FT** |  |
| --- | --- | --- | --- | --- | --- | --- |
| Birth order | 1 |  | 2 |  | 3 |  |
| Weight | 1870g | (3<sup>rd</sup> | 1805g | (3<sup>rd</sup> | 2080g | (10<sup>th</sup> |
|  | percentile) |  | percentile) |  | percentile) |  |
| Height | 44cm | (25<sup>th</sup> | 45cm | (25<sup>th</sup> | 46cm | (50<sup>th</sup> |
|  | percentile) |  | percentile) |  | percentile) |  |
| Cephalic perimeter | 30cm | (50<sup>th</sup> | 30.5cm | (50<sup>th</sup> | 32cm | (50<sup>th</sup> |
|  | percentile) |  | percentile) |  | percentile) |  |
| Apgar score (1') | 9 |  | 8 |  | 7 |  |
| Apgar score (5') | 10 |  | 9 |  | 9 |  |
| Clinical diagnosis at birth | Preterm child<br>Respiratory distress<br>Multifactorial jaundice |  | Preterm child<br>Multifactorial jaundice |  | Preterm child<br>Multifactorial jaundice |  |
| Days at the incubator | 22 |  | 17 |  | 13 |  |

The newborns were kept in an incubator between 2 and 3 weeks. No relevant clinical problems were observed during this period (table 1). Cranial magnetic resonance imaging (MRI) of the two monozygotic twins, performed at age 2;11 (years;months), yielded normal results. No MRI was performed to the fraternal twin. At age 4;3 both monozygotic twins were diagnosed with mild transmission hypoacusia (affecting the left ear in MT1 and both ears in MT2). No hearing problem was observed in the fraternal twin.

### Language and cognitive development

Early developmental milestones were achieved normally by the three children, including head control and the ability to sit without aid and to walk. Nonetheless, their parents and pediatricians soon reported significant developmental differences between the two monozygotic twins and the fraternal twin, including absence of speech, lack of interest in the social environment, and motor disturbances. Differences were prominent at age 7;8, when the global development of the three siblings was compared using the Spanish version of the Battelle Developmental Inventories (figure 1). Below we provide a detailed report of the cognitive and linguistic profiles of the three children.

**Figure 1.**
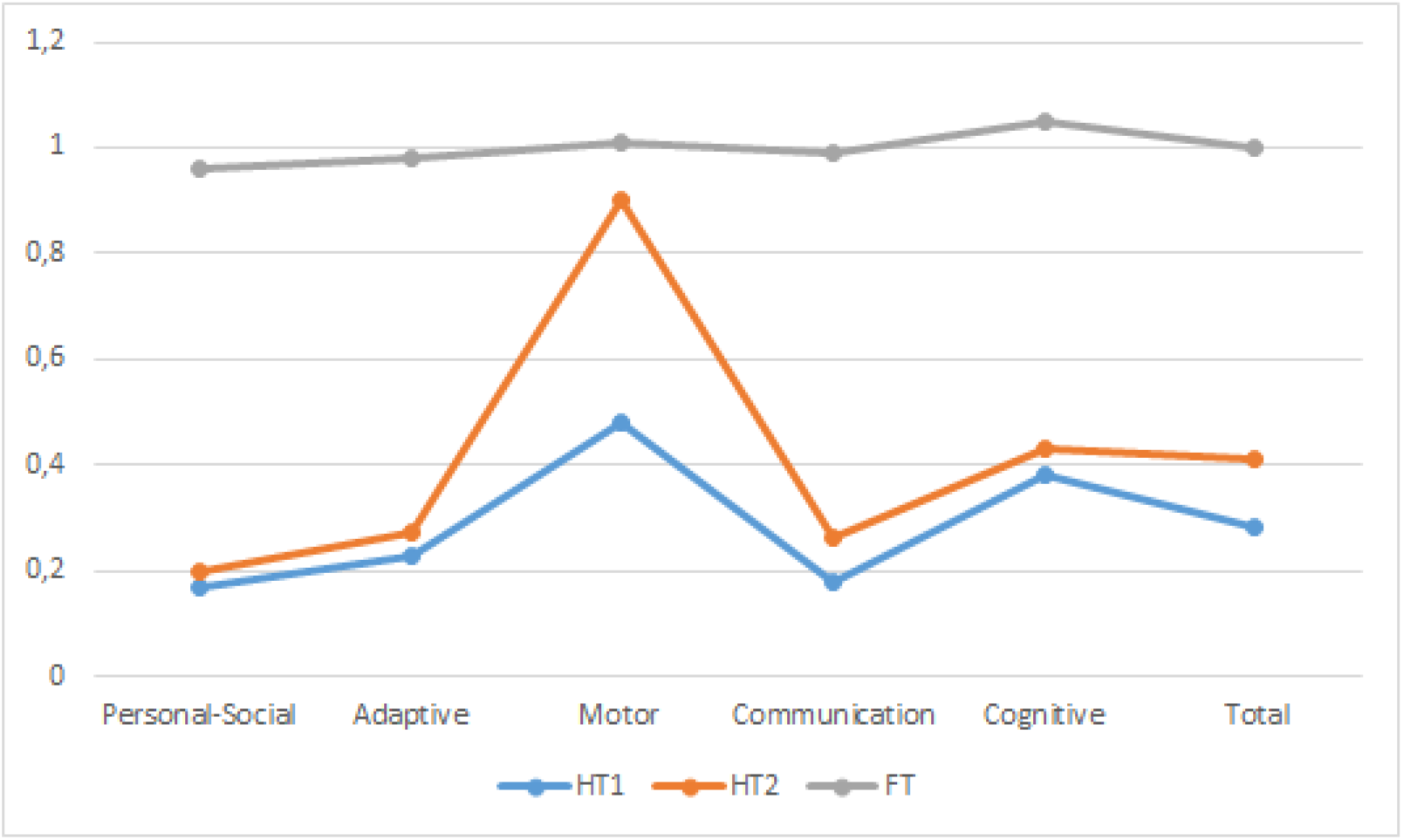
Comparison of the probands’ developmental profiles at age 7;8 according to the Battelle Developmental Inventories. In order to make more reliable comparisons, the resulting scores are shown as relative values referred to the expected scores according to the chronological age of the child. Abbreviations: DA, developmental age; CA, chronological age.

### MZ1 and MZ2

Subsequent developmental stages were achieved later by the two monozygotic twins. At age 2;11 both children exhibited lack of sustained attention and motor disturbances. Language was substantially delayed, to the extent that they only uttered by-syllabic babbling with no evident communicative intention. The two boys only interacted with their closest relatives. They were diagnosed with lack of attention, restlessness, and language delay according to the 9^th^ edition of the International Statistical Classification of Diseases and Related Health Problems (ICD-9).

In order to follow up in detail their global development, the Spanish version of the Battelle Developmental Inventories was administered at ages 3;5, 5;2, and 7;8. The resulting scores were suggestive of a broad developmental delay (which did not ameliorated or exacerbated over the years) impacting mostly on their language abilities. Accordingly, communication skills were severely impaired; cognitive, personal-social, and adaptive abilities were impaired; and motor skills were the less affected area (figures 2A and figures 2B).

**Figure 2.**
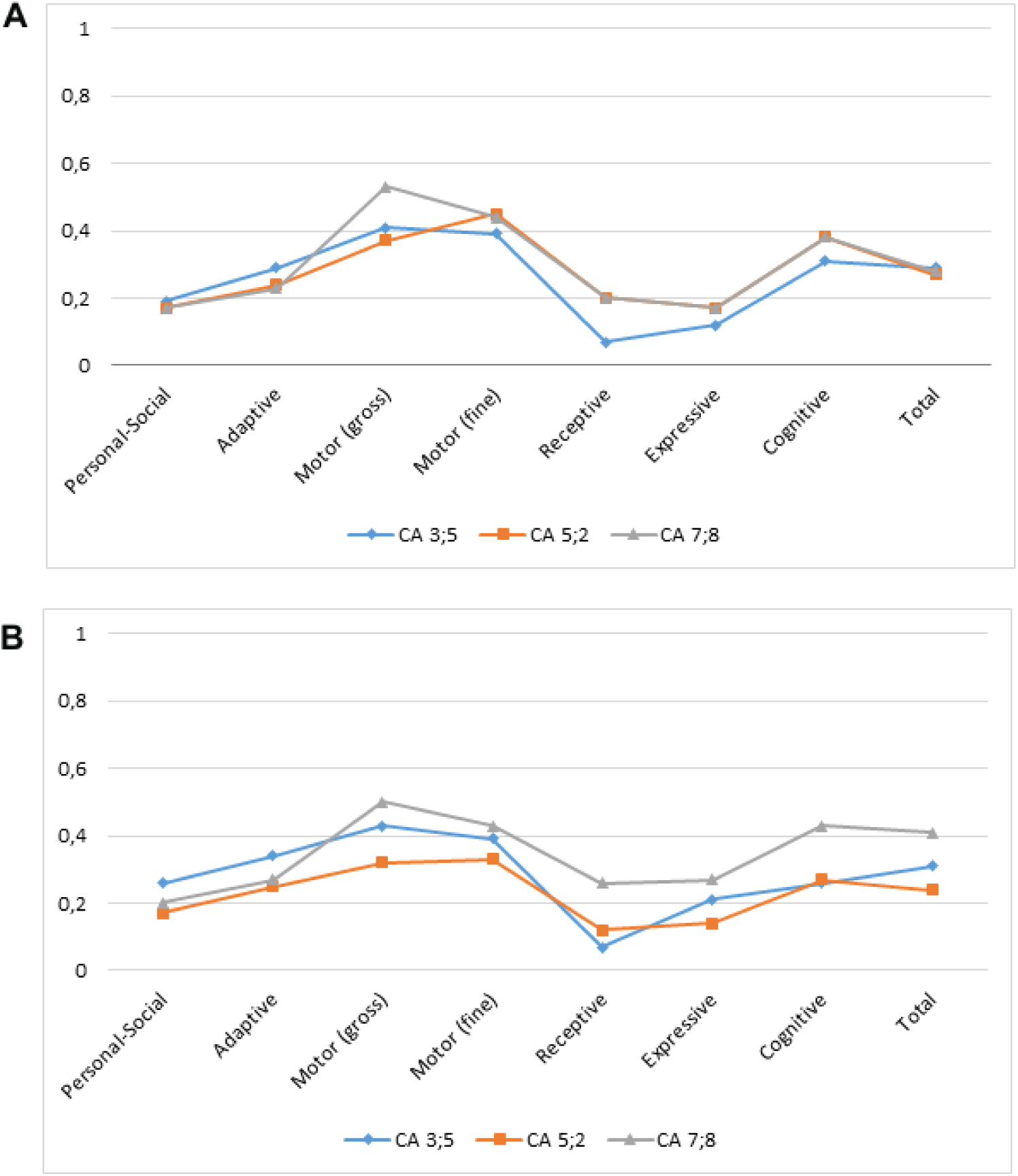
Changes in the developmental profiles of the monozygotic twins from age 3;5 to 7;8 according to the Battelle Developmental Inventories (A: MZ1, B: MZ2). In order to make more reliable comparisons, the resulting scores are shown as relative values referred to the expected scores according to the chronological age of the child. Abbreviations: DA, developmental age; CA, chronological age.

At age 3;5 repetitive behaviour was observed in both children. Although the screening with M-CHAT yielded a positive score, both MT1 and Mt2 were reported to interact normally between them and with their parents, although they didn’t interact with other peers or adults. Gaze following was observed and no obsessive or stereotyped behaviours were reported. In turn, cognitive abilities were found to be severely impaired. Although the kids were able to recognize aspects of their surroundings, like common foods, they were unable to discriminate among colours, sounds, or tactile sensations. Basic concepts like ‘small/big’ or ‘inside/outside’ had not been acquired yet. Language development was seriously delayed. Bi-syllabic babbling was still observed and the children were unable to utter single words. Comprehension was severely impaired, to the extent that they barely understood their own names. Indexical pointing was still absent. Their diagnosis according to the ICD-10 was severe mental delay (F.72). In turn, the children’s motor abilities were quite spared, and they were able to run, jump, throw, and move their arms and legs (still, grasping was absent).

From age 3;2 both children started to attend a general education unit, and also to receive speech therapy 4 times a week following an Applied Behavior Analysis (ABA) paradigm (Baer et al. 1968, Sulzer-Azaroff & Mayer 1991). No major changes at the cognitive, language, and behavioral levels were observed from ages 3 to 4. At age 4, both twins were moved to a special education unit at the general school. At that moment language was still severely delayed. Communicative intentionality was not observed yet. They were unable to attend verbal commands. Speech therapy was also provided at school.

At age 5 stereotyped behaviour and restricted interests were observed, supporting the presence of autism. Auto-stimulatory behaviour was also reported, as well as lack of sustained attention, and motor hyperactivity. Regarding their communicative abilities, auxiliary gestures were not observed and both children were still unable to understand verbal cues or instructions. Their diagnosis according to the ICD-10 was ASD (F.84.1) and severe mental delay (F.72).

At age 6 intentionality was observed in some verbal interactions and both children seemed to understand simple verbal commands. Deictical indexing was also reported. In order to improve their verbal abilities, the Picture Exchange Communication System (PECS) (Brondy and Frost 1998) was introduced by the speech therapist at school. This is an augmentative system intended to provide the child with a self-initiating, functional communication system. The system makes use of simple icons that are arranged in “sentence” structures, with a focus on requesting instead on responding or commenting. Both children started to use single words for requesting, although some differences between HT1 and HT2 were observed. Accordingly, HT1 only completed the phase 2 of PECS (Expanding Spontaneity), while HT2 was able to complete the phase 3 (Picture Discrimination). Some differences were observed in their behaviour too. Accordingly, HT1 was more defiant, but less aggressive than HT2; self-hurting was reported in HT2 only.

From age 7 some regression was observed in both children at the behavioral level, with an impact on their language abilities. Stereotypies and restricted interests, and aggressive behaviour was reported of HT1, whereas inattention problems were observed in HT2. At present both twins have been moved to a specialised school for disabled children.

### Fraternal twin

Early in childhood parents reported language delay, particularly in the expressive domain, and the child received speech therapy until age 4 to improve verbal expression, and speech fluency and articulation. At age 3 he commenced attending a general school. At that moment his speech was impaired and he suffered from articulatory problems and apraxia. The child employed an unintelligible jargon with their peers and seemed unable to understand simple commands, which suggests that he suffered from some deficit in comprehension too. Nonetheless, these problems disappeared with time and evidence of language delay was not observed from age 5 onwards. Also, no delays or deficits were reported regarding his broad cognitive development and the achievement of instrumental abilities. In order to evaluate his global development in detail, the Spanish version of the Battelle Developmental Inventories was administered at age 7;8. The proband scored like typically-developing children in all domains (Figure 3A). Psycholinguistic development was assessed in more detail with the Spanish version of the Illinois Test of Psycholinguistic Abilities (ITPA) and with the verbal component of the Spanish version of the WISC-IV. In the ITPA the child scored 306, showing a composite psycholinguistic age of 9;1, well above his chronological age. However, his profile was quite irregular, with marked strengths and weakness, particularly in the automatic domain, in which he scored the lowest in the grammar closure test, which evaluates the ability to add missing grammatical components to a given sentence (figure 3B). In the WISC-IV the proband scored 10 in the Verbal Comprehension Index (VCI), with lower scores in Similarities and higher scores in Comprehension and Information (figure 3C). His global score was similar to the scores achieved by his parents in the Intellectual Verbal Index of the WAIS-III, although the latter scored lower at the Comprehension and the Information components. All of them scored similarly in the tasks evaluating working memory (figure 3C).

**Figure 3.**
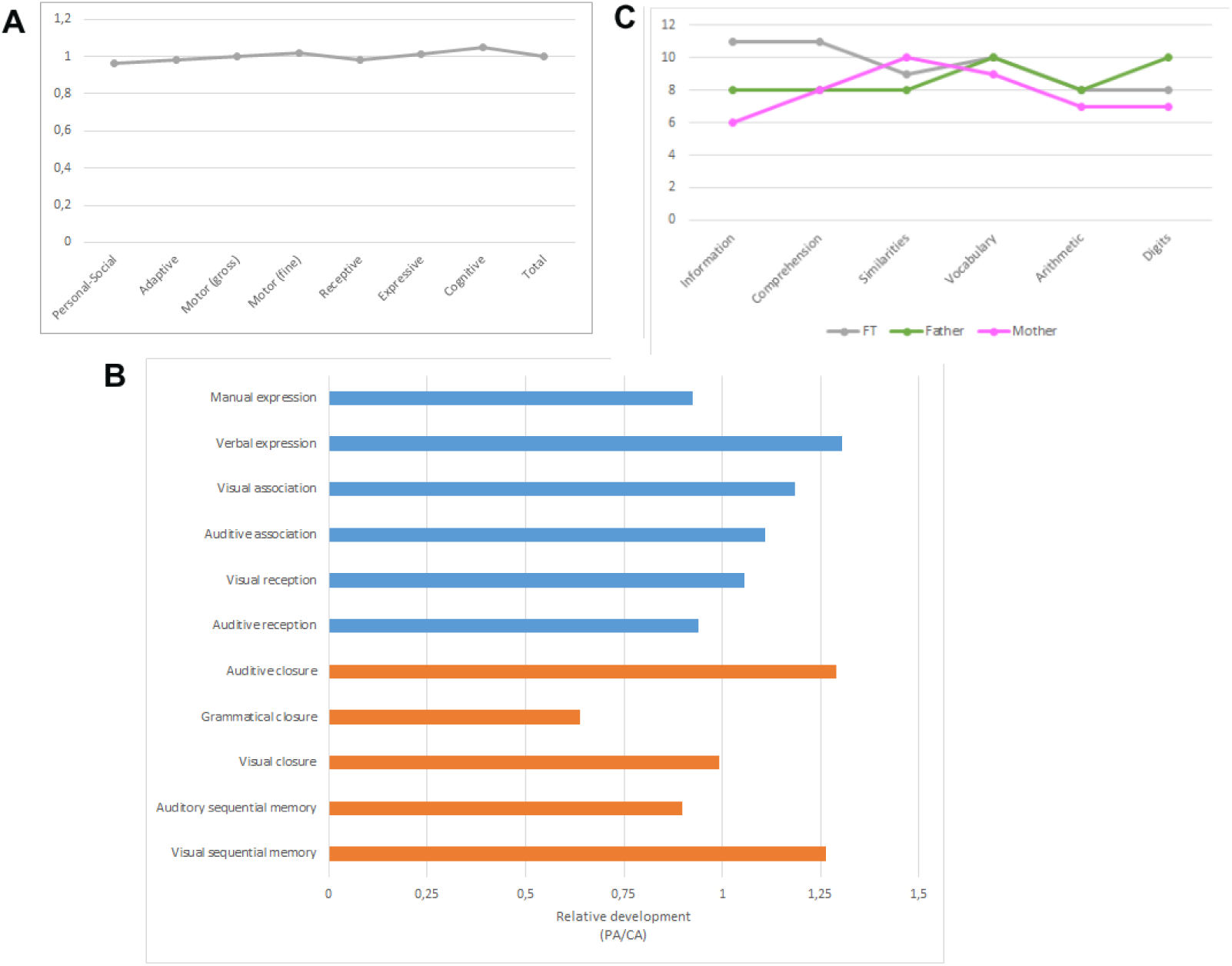
Developmental profile and language abilities of the fraternal twin at age 7;8. A. Developmental profile according to the Battelle Developmental Inventories. B. Developmental profile according to the ITPA. C. Developmental profile according to the verbal component of the WISC-IV (for comparison, the figure includes the parents’ profiles according to the verbal component of the WAIS-III).

### Molecular Cytogenetic Analysis

Routine molecular cytogenetic analyses of the three children were performed at age 3. FISH analyses of the loci of *SNRPN* and *UBE3A* (two main determinants of Angelman’s syndrome) were normal. No major chromosomal rearrangements were observed in the karyotype analysis. At age 5;7 a MLPA of the subtelomeric regions of all chromosomes was performed. A deletion in 4q35.2 affecting *FRG1* was then reported, but this was not confirmed later by FISH when using a 4qter subtelomeric probe. In order to discard the presence of low-frequency mosaicisms and/or cryptic chromosomal alterations, a comparative genomic hybridization array (array-CGH) was performed. The array discarded the deletion in chromosome 4, but identified in the three children a duplication in the 15q11.2 region that was inherited from the father (figure 4A and table 2). Differences in the predicted size of the duplicated fragments were expected to result from differences in the intensity of the signal, considering that the four probes exhibited similar profiles. The duplicated region corresponded to the BP1-BP2 region at 15q11.2, which encompasses the genes *TUBGCP5*, *CYFIP1*, *NIPA2*, and *NIPA1*. The three children also bear a microduplication in 6p25.3 which affects the promoter and part of the first exon of *DUSP22* and which was inherited from the mother (figure 4B and table 2).

**Figure 4.**
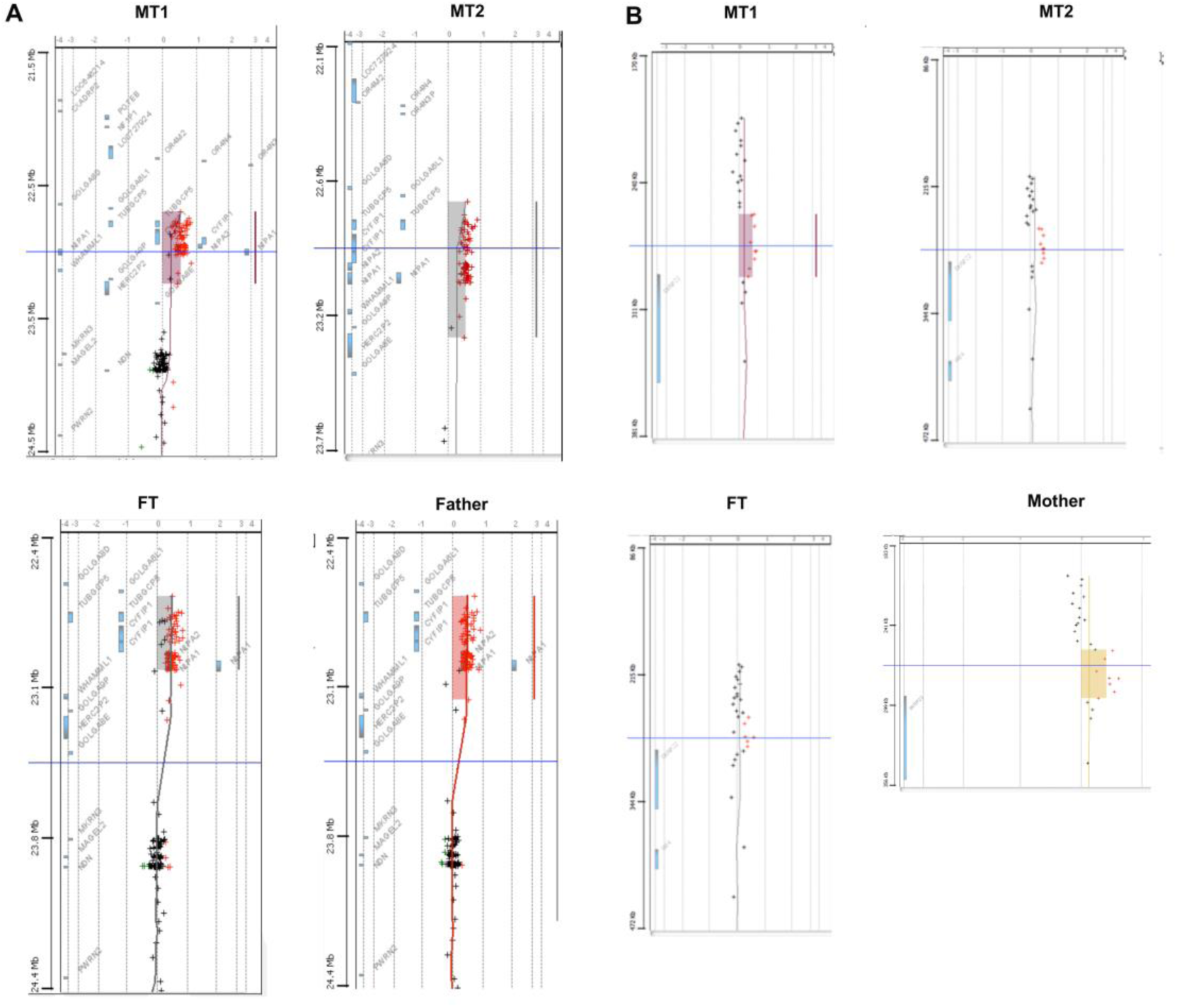
Array-CGH and duplicated genes in the probands. A. Array-CGH of the probands’ chromosome 15 showing the microduplication at 15q11.2. B. Array-CGH of the probands’ chromosome 6 showing the microduplication at 6p25.3.

**Table 2.**
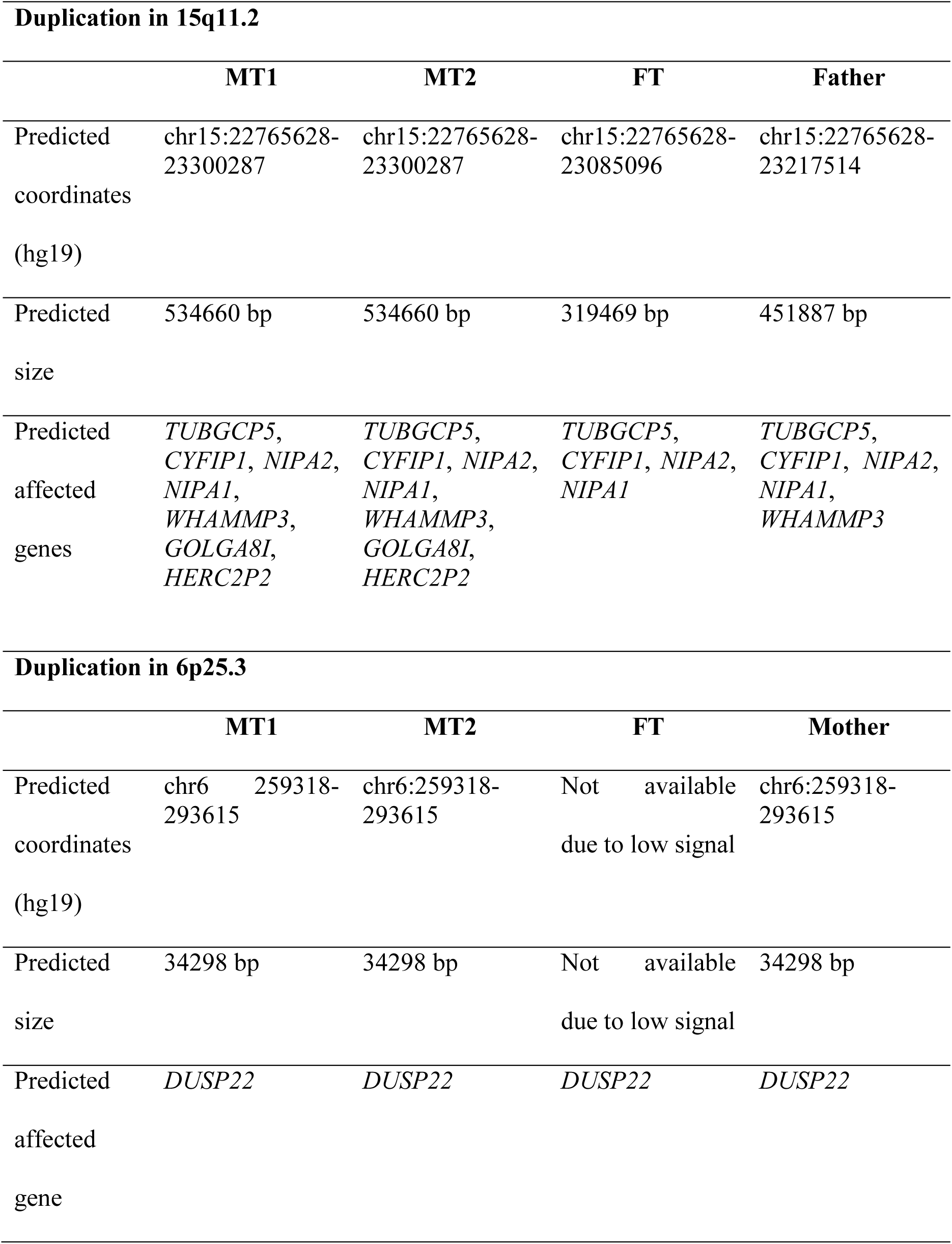
Summary table with the CNVs found in the probands.

## Discussion

In this paper we have characterized in detail the linguistic and cognitive profile of a family (the father and three male siblings) bearing a duplication of the BP1-BP2 region at 15q11.2. Two of the children are monozygotic twins who exhibit most of the features commonly associated to this duplication (table 3). The father and the fraternal twin are phenotypically normal carriers, although the fraternal twin exhibited some language delay in early infancy which resolved with time. This outcome can be explained in terms of reduced penetrance or altered gene dosage on a particular genetic background, as reported in other cases (Burnside et al. 2011).

**Table 3.**
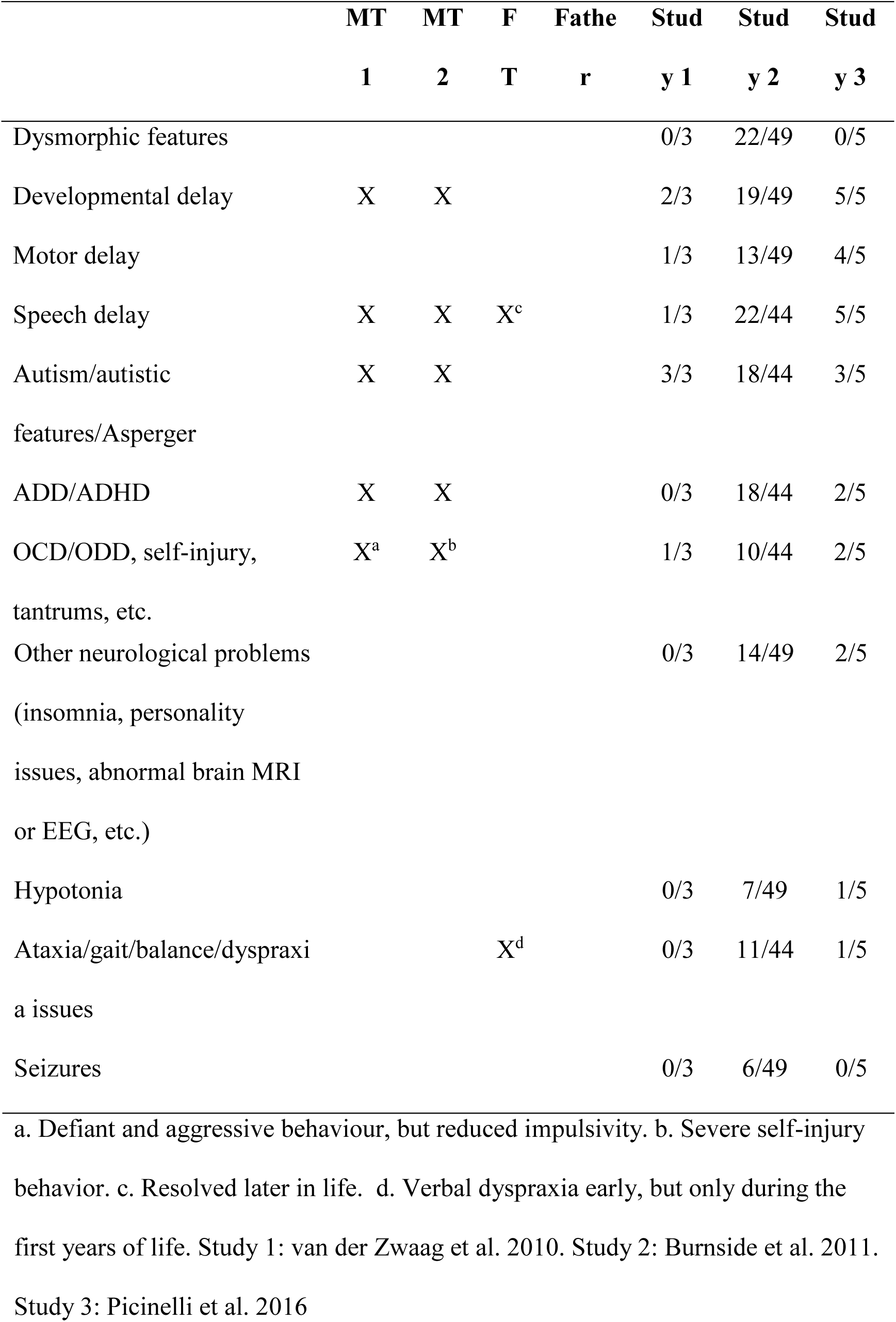
Summary table with the clinical findings of the probands compared to those described in other patients bearing the duplication of the BP1-BP2 region

The core BP1-BP2 region contains four genes (*TUBGCP5*, *CYFIP1*, *NIPA2*, and *NIPA1*) involved in behavioral and neurological function, which are expressed biallelically (Chai et al. 2003). As reviewed by Picinelli et al. (2016), *TUBGCP5* has been related to Attention Deficit-Hyperactivity Disorder (ADHD) and Obsessive-Compulsive Disorder (OCD); *CYFIP1* encodes a protein that interacts with FMRP, the main determinant for Fragile X Mental Retardation syndrome; and *NIPA1* has been linked to autosomal dominant hereditary spastic paraplegia. No patient bearing a single duplication of any of these genes is found in DECIPHER. Nonetheless, stronger links between some of these genes and language (dis)abilities are expected to exist. Accordingly, a common variant of *CYFIP1* has been recently associated with inter-individual variation in surface area across the left supramarginal gyrus, a brain area involved in language processing, and the gene is predicted to be regulated by *FOXP2*, a well-known gene important for speech and language (Woo et al. 2016).

The three children in our study also bear a partial duplication of the gene *DUSP22*, involving the proximal part of the promoter and the first exon, which is also found in their mother. This duplication has been reported as common polymorphism in the normal population, but might underlie differential susceptibility to disease (Iafrate et al. 2004, suppl. table 1). *DUSP22* encodes a dual specificity phosphatase involved in cell death (Ju et al. 2016). Similar duplications are reported as of unknown pathogenicity in DECIPHER (patient 250238), whereas deletions affecting the same region are reported as likely benign (patient 286128) or of unknown pathogenicity (patient 250711). CNVs affecting the promoter of the gene only have been found in individuals with intellectual disability (DECIPHER patients 1566 (duplication), and 652 (deletion)). Interestingly, the promoter of *DUSP22* has been found hypermethylated in the hippocampus of patients with Alzheimer’s disease and the DUSP22 protein seemingly contributes to determine TAU phosphorylation status and CREB signaling (Sánchez-Mut et al. 2014). DECIPHER database shows that duplications of the whole gene *DUSP22* gene are less frequently found than deletions, which usually entail more severe developmental problems, including autism (Leblond et al. 2012). Speech and language delay is only reported in patients with deletions affecting the downstream genes (DECIPHER patients 249742 or 273907). Because of the normal phenotype of the mother, we expect that the deficits observed in the two affected children result from the duplication of the BP1-BP2 region at 15q11.2. Nonetheless, we cannot rule out the possibility that a change in the amount of DUSP22 due to the partial duplication of the gene contributes to the language deficits found in the three siblings at the initial stages of development, considering that the mother scored below the mean in the tasks evaluating verbal comprehension.

## CONCLUSIONS

Although the exact genetic cause of the language and cognitive impairment exhibited by two of our probands remains to be fully elucidated, we believe that the duplication of the genes at 15q11.2 discussed above may explain most of their deficits and clinical problems. Comparisons with the genetic background of their unaffected brother and father will help clarify the variable manifestation of the duplication of this region. We expect that this familiar case contribute as well to a better understanding of the genetic underpinnings of the human faculty for language.

## ETHICS, CONSENT, AND PERMISSIONS

Ethics approval for this research was granted by the Comité Ético del Hospital Universitario “Reina Sofía”. Written informed consent was obtained from the probands’ parents for publication of this case report and of any accompanying tables and images. A copy of the written consent is available for review by the Editor-in-Chief of this journal.

## LIST OF ABBREVIATIONS

ADD: Attention Deficit Disorder
ADHD: Attention Deficit-Hyperactivity Disorder
CA: chronological age.
CGH: comparative genomic hybridization
CNVs: copy number variants
DA: developmental age
FISH: fluorescence in situ hybridization
ICD: International Statistical Classification of Diseases and Related Health Problems
ITPA: Illinois Test of Psycholinguistic Abilities
M-CHAT: Modified Checklist for Autism in Toddlers
MLPA: Multiplex Ligation-dependent Probe Amplification
MRI: magnetic resonance imaging
OCD: Obsessive-Compulsive Disorder
ODD: Oppositional Defiant disorder
PCR: polymerase chain reaction
PECS: Picture Exchange Communication System
WAIS: Wechsler Adult Intelligence Scale
WISC: Wechsler Intelligence Scale for Children

## AUTHORS’ CONTRIBUTIONS

ABB interpreted the data and wrote the paper. MBM and IEP performed the molecular cytogenetic analyses. MJR assessed the cognitive and linguistic problems of the child, and made substantial contributions to the analysis of the data. All authors revised the draft of the paper, and read and approved the final manuscript.

## AUTHORS’ INFORMATION

Antonio Benítez-Burraco is a Research Fellow at the Modern Languages Faculty of the University of Oxford and Associate Professor at the Department of Philology of the University of Huelva (Spain). Monserrat Barcos Martínez and Isabel Espejo Portero are staff physicians at the Laboratory of Molecular Genetics of the University Hospital “Reina Sofía” in Córdoba (Spain) and members of the Maimónides Institute of Biomedical Research at Córdoba (Spain). M^a^ Salud Jiménez Romero is an Assistant Professor at the Department of Psychology of the University of Córdoba (Spain) and also a member of the Maimónides Institute of Biomedical Research at Córdoba (Spain)

## ACKNOWLEDGMENTS

We would like to thank the probands and their family for their participation in this research. We wish thank also Luis Antonio Alcaraz Mas, from Bioarray (http://www.bioarray.es/es), for his technical aid with the microarray analyses. Preparation of this work was supported in part by funds from the Spanish Ministry of Economy and Competitiveness (grants FFI2014-61888-EXP and FFI2016-78034-C2-2-P to Antonio Benítez-Burraco).

